# Active cortical networks promote shunting fast synaptic inhibition *in vivo*

**DOI:** 10.1101/2023.03.01.530641

**Authors:** Richard J. Burman, Paul J.N. Brodersen, Joseph V. Raimondo, Arjune Sen, Colin J. Akerman

## Abstract

Fast synaptic inhibition determines neuronal response properties in the mammalian brain and is mediated by chloride-permeable ionotropic GABA-A receptors (GABA_A_Rs). Despite their fundamental role, it is still not known how GABA_A_Rs signal in the intact brain. Here we use *in vivo* gramicidin recordings to investigate synaptic GABA_A_R signaling in mouse cortical pyramidal neurons under conditions that preserve native transmembrane chloride gradients. In anaesthetized cortex, synaptic GABA_A_Rs exert classic hyperpolarizing effects. In contrast, GABA_A_R-mediated synaptic signaling in awake cortex is found to be predominantly shunting. This is due to more depolarized GABA_A_R equilibrium potentials (E_GABAAR_), which are shown to result from the high levels of synaptic activity that characterize awake cortical networks. The E_GABAAR_ observed in awake cortex can facilitate the decoupling of local networks, which improves the ability of the network to discriminate stimuli. Our findings therefore suggest that GABA_A_R signaling adapts to optimize cortical functions.

## Introduction

Synaptic inhibition is tightly coupled to synaptic excitation and plays a key role in cortical computations^1^, including modulating sensory response properties^2,3^ and oscillatory activities^4^. Fast synaptic inhibition in cortex is mediated by ionotropic GABA-A receptors (GABAAR), which are primarily permeable to chloride (Cl^-^)^5,6^. The inhibitory effects that GABAARs have upon a neuron, therefore, depend upon the local transmembrane Cl^-^ gradient, which reflects a dynamic equilibrium between Cl^-^ extrusion and intrusion processes^7–9^.

When the GABAAR equilibrium potential (EGABAAR) is more negative than the membrane potential (V_m_), GABA_A_R activation will lead to membrane hyperpolarization. If E_GABAAR_ is close to V_m_, GABA_A_R activation would have minimal effects upon V_m_ and the inhibitory effects would be primarily mediated by local effects on the membrane resistance (R_m_) – a phenomenon known as “shunting inhibition”^10,11^. These different forms of signaling determine how the inhibition is temporally integrated with incoming excitatory synaptic inputs^12,13^. Thus, appreciating how fast synaptic inhibition operates is fundamental to understanding how neuronal and network activity are regulated in the intact brain.

Previous work has identified the contribution of different transmembrane Cl^-^ fluxes in determining synaptic E_GABAAR_ in cortex. This includes Cl^-^ effluxes mediated by the potassium-chloride cotransporter, KCC2^14,15^, as well as Cl^-^ influxes such as those mediated by GABA_A_Rs themselves, which can vary depending upon a neuron’s activity^16,17^. Previous investigations of synaptic E_GABAAR_ have relied heavily upon *in vitro* investigation and therefore been influenced by the distorted fluxes that operate under these recording conditions^8^. Consequently, there is a lack of evidence regarding how fast synaptic inhibition operates in the intact cortex.

To address this gap, we combine optogenetic activation of GABAergic synaptic inputs with *in vivo* gramicidin perforated patch-clamp recordings. This enables us to measure synaptic E_GABAAR_ and GABA_A_R driving forces in the intact rodent cortex, under conditions in which transmembrane Cl^-^ gradients are preserved. We demonstrate that in contrast to the anaesthetized cortex, pyramidal neurons of the awake cortex exhibit a relatively high synaptic E_GABAAR_, which is close to resting V_m_ and generates a clear preference for shunting fast synaptic inhibition. This depolarized E_GABAAR_ results from the high levels of synaptic activity in the awake cortex, and computational modelling indicates that this facilitates the decoupling of active cortical networks to improve stimulus discrimination.

## Results

### Measuring the equilibrium potential and driving force for synaptic GABA_A_Rs *in vivo*

To study GABA_A_R-mediated synaptic signaling *in vivo*, we established gramicidin perforated patch-clamp recordings from layer 2/3 (L2/3) pyramidal neurons in primary visual cortex (V1) of anaesthetized mice aged six to eight weeks (**Figure 1A**). Similar to *in vitro* findings^18,19^, the perforation of the neuronal membrane with cation-selective gramicidin pores was marked by a decrease in the series resistance (R_s_; **Figure 1B**). During the ten minutes following gigaseal formation we observed a 20-fold decrease in R_s_ (mean R_s-0min_: 1126.9 ± 35.8 MΩ vs. R_s-10min_: 50.2 ± 3.8 MΩ), which provided stable conditions for studying synaptic responses. For our experiments, we used transgenic mice expressing channelrhodopsin-2 (ChR2) in Gad2-positive neurons (**Figure 1A** and **1C**). This enabled us to initiate presynaptic GABA release by optically activating the axons of nearby GABAergic interneurons^20,21^, resulting in postsynaptic GABAARs responses in the L2/3 pyramidal neuron that could be recorded in either current clamp or voltage clamp (**Figure 1D**).

**Figure 1:**
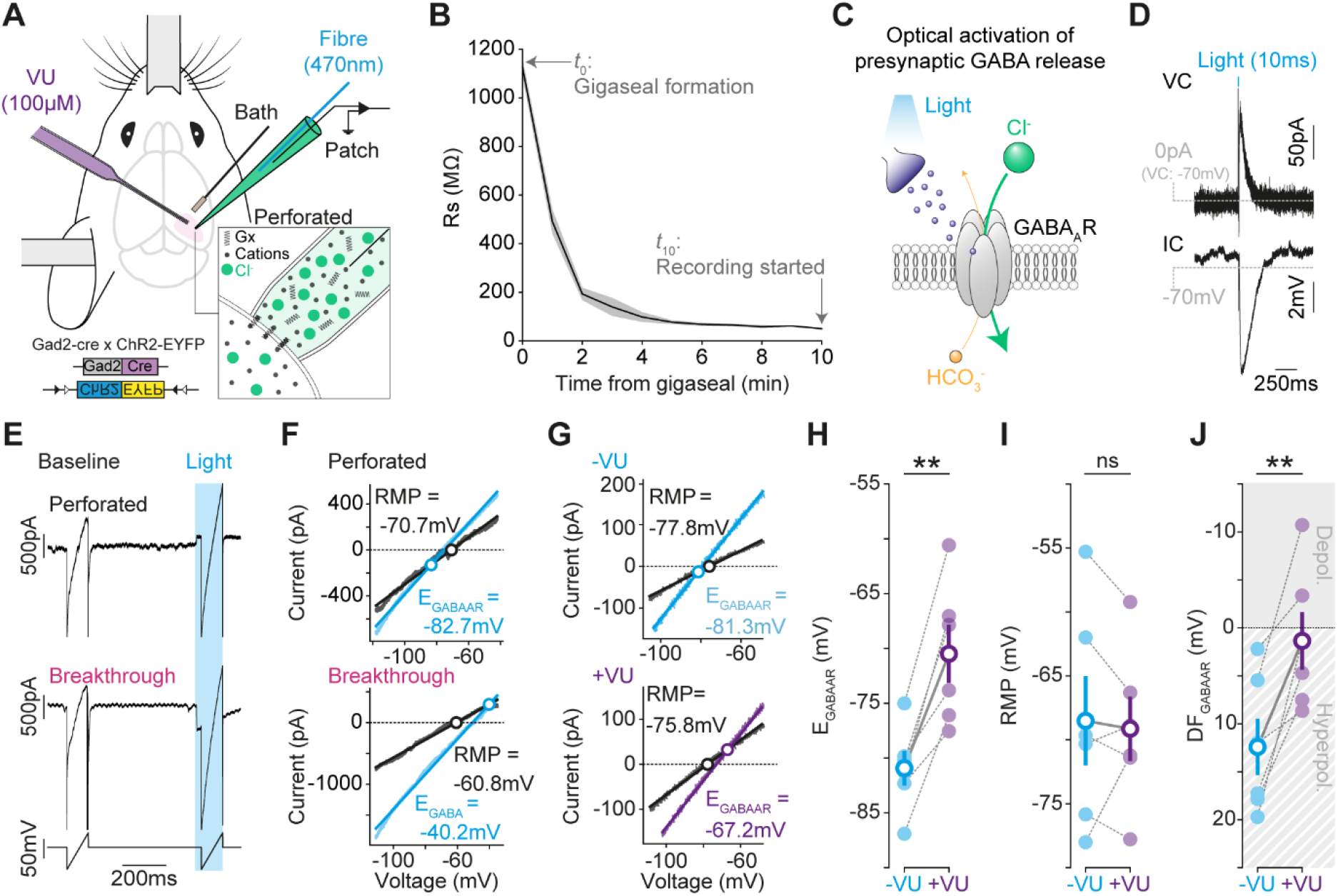
Measuring synaptic E_GABAAR_ and synaptic GABA_A_R driving forces *in vivo*. **(A)** Setup for performing gramicidin perforated patch-clamp recordings in V1, in combination with optogenetic activation of local GABAergic synaptic inputs. Mice expressed ChR2-EYFP in Gad2-positive interneurons. Inset illustrates gramicidin creating cation-selective pores in the neuronal membrane, which preserve intracellular Cl^-^. Light pulses were delivered via an optic fiber within the patch pipette. In some experiments, a second pipette was used to deliver the KCC2 antagonist, VU0463271 (VU). (**B**) Series resistance (R_s_) over time (mean ± SEM; *n* = 10 neurons, 7 mice). (**C**) Optogenetic approach elicits presynaptic GABA release and activates postsynaptic GABA_A_Rs. (**D**) Light-evoked currents (VC) and potentials (IC). (**E**) Ramp protocol in perforated configuration (top) and following breakthrough into whole-cell configuration (middle). The voltage protocol before R_s_ correction is also shown (bottom) and consisted of a control ramp (‘baseline’) and a second ramp during the light-evoked synaptic GABA conductance (‘light’). (**F**) IV plots for baseline (black) and light (cyan) ramps performed under perforated (top) and breakthrough (bottom) conditions. The reversal potential of the baseline current (‘RMP’) and EGABAAR are indicated with circles. (**G**) IV plots showing synaptic E_GABAAR_ before (-VU, cyan, top) and after VU application (+VU, purple, bottom). (**H**) Population data (*n* = 6 neurons from 6 mice) showed a depolarizing shift in synaptic E_GABAAR_ after VU (-VU: -80.9 ± 1.6 mV vs. +VU: -70.9 ± 2.6 mV; *p* = 0.002, *paired t-test*). (**I**) VU did not affect RMP (-VU: -68.5 ± 3.5 mV vs. +VU: -69.1 ± 2.5 mV; *p* = 0.73, *paired t-test*). (**J**) VU caused a depolarizing shift in GABA_A_R driving force (DF_GABAAR_ = RMP - E_GABAAR_, -VU: 12.4 ± 2.9 mV vs. +VU: 1.4 ± 3.0 mV; *p* = 0.006, *paired t-test*). ‘ns’, non-significant; **, *p* < 0.01.

To measure synaptic E_GABAAR_ *in vivo*, we combined our optogenetic approach with short voltage-ramp protocols (**Figure 1E**) to minimize disruption to transmembrane Cl^-^ gradients and generate current-voltage (IV) plots from which the resting membrane potential (RMP) and E_GABAAR_ can be determined (**Figure 1F**). To establish the integrity of the perforated patch-clamp recordings, the internal pipette solution contained a high concentration of Cl^-^ (∼150 mM) so that the status of the patch at the end of a recording could be confirmed by electing to breakthrough into whole-cell mode (**Figure 1E** and **1F**).

To check that our voltage-ramp protocols provided an accurate estimate of synaptic E_GABAAR_, results were compared to voltage step protocols performed in the same neuron (**Figure S1A-D**). Step protocols take longer to perform, but offer the chance to analyze synaptic E_GABAAR_ at a defined time following presynaptic GABA release^22^, thereby further isolating the GABA_A_R response (**Figure S1A-D**). The estimates of synaptic E_GABAAR_ from ramp protocols were found to be equivalent to those made using step protocols (**Figure S1E**). The synaptic E_GABAAR_ measurements from the ramp protocols were not related to the amplitude of the synaptic GABA_A_R response (**Figure S2A**), or the neuron’s R_s_ (**Figure S2B**).

To further validate the setup, we assessed whether our *in vivo* measurements were sensitive to changes in synaptic E_GABAAR_ that are caused by altering the balance of transmembrane Cl^-^ fluxes. As expected, application of the selective KCC2 antagonist, VU0463271 (VU), resulted in a depolarizing shift in synaptic E_GABAAR_ (**Figure 1G** and **1H**), consistent with the blocking of a Cl^-^ efflux. Whilst VU caused a depolarizing shift in synaptic E_GABAAR_, there was no detectable change in RMP (**Figure 1I**), and consequently a depolarizing shift in the driving force for synaptic GABA_A_Rs (DF_GABAAR_ = RMP – E_GABAAR_; **Figure 1J**).

### Awake cortex exhibits more depolarized synaptic E_GABAAR_ and shunting fast synaptic inhibition

Having established gramicidin recordings *in vivo*, we investigated synaptic GABA_A_R signaling in the awake cortex (**Figure 2A**). In keeping with previous whole-cell recording studies, our gramicidin recordings in head-fixed mice revealed that L2/3 pyramidal neurons in the awake cortex exhibit high levels of synaptic activity. Compared to our recordings in anaesthetized cortex, awake neurons exhibited more depolarized membrane potentials (V_m_, **Figure 2B** and **2C**) and larger fluctuations in their subthreshold V_m_ (**Figure 2D**). This is consistent with the elevated levels of excitatory and inhibitory synaptic activity reported in the awake cortex^23,24^. The membrane resistance (R_m_) was also lower in the awake recordings (An.: 87.2 ± 6.1 MΩ, *n* = 14 neurons, 8 mice vs. Aw.: 50.6 ± 6.4 MΩ, *n* = 13 neurons, 10 mice, *p* < 0.0001, *unpaired t-test*), consistent with the increased membrane conductance associated with high levels of synaptic activity^25^.

**Figure 2:**
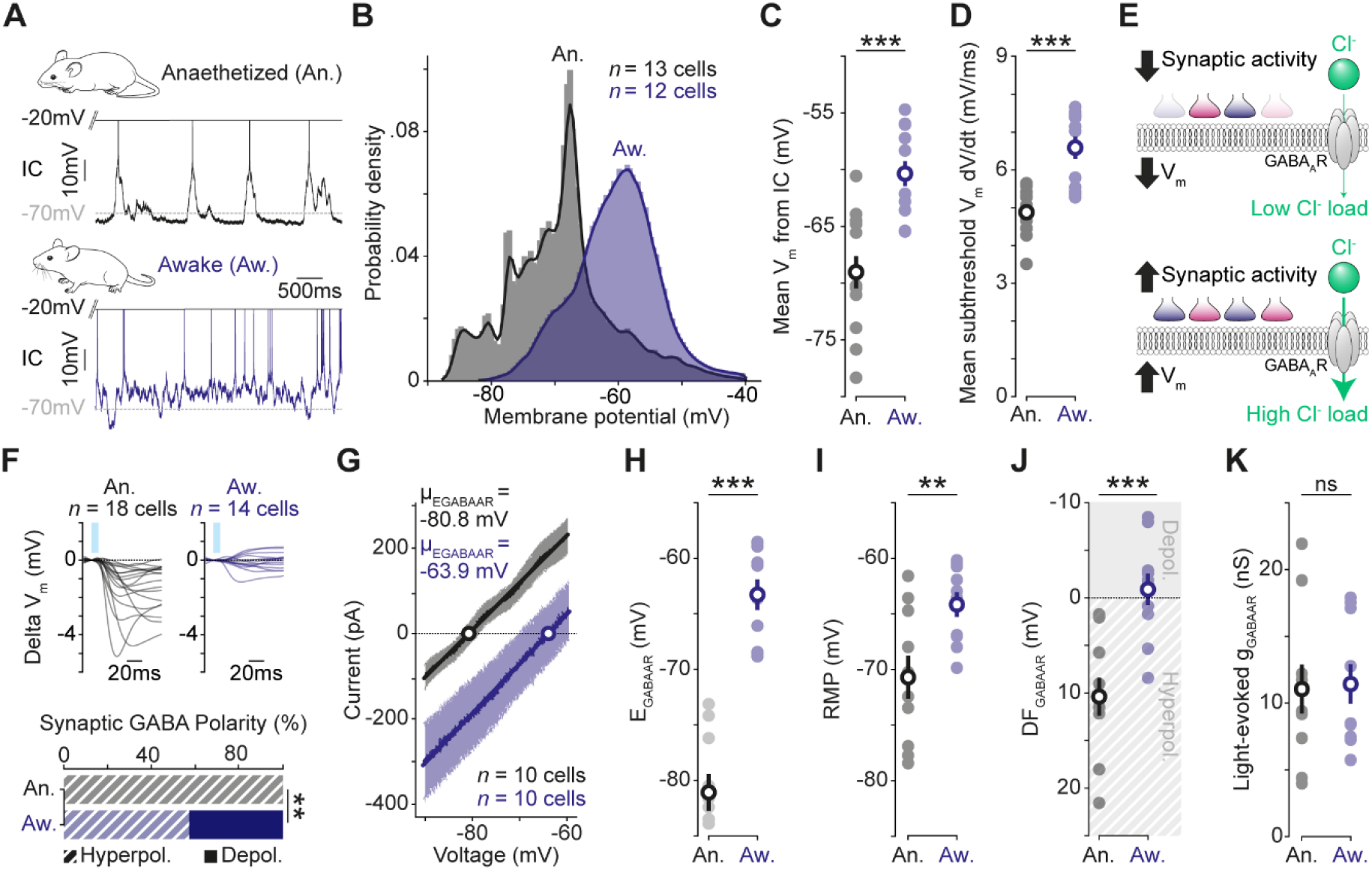
Awake cortex exhibits depolarized synaptic EGABAAR and shunting fast synaptic inhibition. **(A)** Current-clamp (IC) recording of spontaneous activity from a L2/3 pyramidal neuron in an anaesthetized (black, top) and awake mouse (blue, bottom). (**B**) Probability density function for V_m_ in anaesthetized (*n* = 13 cells, 7 mice) and awake cortex (*n* = 12 cells, 10 mice). (**C**) Mean V_m_ was more depolarized in awake cortex (An.: -69.1 ± 1.4 mV vs. Aw.: -60.3 ± 1.1 mV; *p* < 0.0001, *unpaired t-test*). (**D**) Mean change in subthreshold V_m_ was greater in awake cortex (An.: 4.9 ± 0.2 mV/ms vs. Aw.: 6.6 ± 0.3 mV/ms; *p* < 0.0001, *unpaired t-test*). (**E**) Illustration of how a more depolarized V_m_ and higher synaptic activity are conducive to greater GABA_A_R-mediated Cl^-^ influxes. (**F**) Averaged light-evoked postsynaptic IC responses in anaesthetized (*n* = 18 cells, 10 mice) and awake cortex (*n* = 14 cells, 11 mice). Responses in awake cortex produced V_m_ deflections that remained close to the RMP and could be depolarizing or hyperpolarizing (An.: Depol. 0/18 vs. Aw.: Depol. 6/14; *p* = 0.003, *Fisher-Exact test*). (**G**) Summary IV plot of all anaesthetized (*n* = 10 cells, 6 mice) and awake (*n* = 10 cells, 9 mice) light-evoked GABA currents reveal a more depolarized synaptic E_GABAAR_ in awake cortex. (**H**) Synaptic E_GABAAR_ is more depolarized in awake cortex (An.: -81.1 ± 1.7 mV vs. Aw.: -63.3 ± 1.4 mV; *p* < 0.0001, *unpaired t-test*). (**J**) VC recordings confirmed a more depolarized RMP in awake cortex (An.: -70.7 ± 1.9 mV vs. Aw.: -64.2 ± 1.1 mV; *p* = 0.008, *unpaired t-test*). (**K**) GABAAR driving force (DF_GABAAR_ = RMP – E_GABAAR_) was more depolarized in awake cortex (An.: 10.4 ± 2.0 mV vs. Aw.: -0.9 ± 1.7 mV; *p* = 0.0004, *unpaired t-test*). Awake DF_GABAAR_ was not different to zero, consistent with synaptic GABA_A_Rs exerting a predominantly shunting effect (An.: *p* = 0.0006, *one-sample t-test*; Aw.; *p* = 0.61, *one-sample t-test*). ‘ns’, non-significant; **, *p* < 0.01; ***, *p* < 0.001.

The high level of synaptic activity in the awake cortex includes strong activation of GABA_A_R-containing synapses^3^. This generates conditions that are predicted to increase the Cl^-^ influxes (i.e. a ‘Cl^-^ load’) experienced by a cortical neuron^8,26^, particularly as more depolarized V_m_ values will increase the driving force for Cl^-^ to enter via the GABA_A_R (**Figure 2E**). We hypothesized that the conditions of the awake cortex would impact transmembrane Cl^-^ gradients, and thereby affect synaptic GABA_A_R signaling. To test this prediction, we used our optogenetic approach to investigate synaptic GABA_A_R responses, comparing these between the anaesthetized and awake cortex.

Firstly, current-clamp recordings revealed that light-evoked synaptic GABA_A_R responses in the awake cortex produced V_m_ deflections that remained close to the RMP and could be depolarizing or hyperpolarizing (**Figure 2F**). This indicated a net shunting effect for synaptic GABA_A_R in the awake cortex, consistent with a different DF_GABAAR_ that could be caused by a more depolarized synaptic E_GABAAR_. We confirmed this by performing ramp protocols in voltage clamp, demonstrating that synaptic E_GABAAR_ was more depolarized in the awake cortex (**Figure 2G** and **2H**). Whilst the RMP was also more depolarized in the awake cortex (**Figure 2I**), the net effect was to move the synaptic DF_GABAAR_ in a depolarized direction when compared to anaesthetized cortex (**Figure 2J**). Importantly, the synaptic DF_GABAAR_ in the awake cortex was not different from zero, consistent with the conclusion that GABA_A_R-mediated synaptic signaling favors shunting inhibitory effects in the awake state. These differences were not related to how the synaptic GABA_A_Rs were activated, because the amplitude of the light-evoked synaptic GABA_A_R conductances were comparable in the anaesthetized and awake cortex (**Figure 2K**). Meanwhile modelling of synaptic GABA_A_R responses in biologically realistic neurons confirmed that differences in synaptic E_GABAAR_ could not be explained by the potential effects of intrinsic membrane properties between anaesthetized and awake cortex (**Figure S3**).

### Network activity in awake cortex raises synaptic EGABAAR and promotes shunting fast synaptic inhibition

Having established that awake cortex favors shunting synaptic GABA_A_R responses, we explored whether the more depolarized synaptic E_GABAAR_ is caused by the high levels of synaptic activity that exist in the awake state. To test this, we examined the effects of reducing local network activity in awake cortex by blocking glutamatergic signaling with a local injection of the AMPA receptor antagonist, NBQX (**Figure 3A**). Following NBQX injection, the distribution and mean V_m_ of cortical neurons was more hyperpolarized (**Figure 3B** and **3C**), and fluctuations in subthreshold V_m_ were greatly decreased (**Figure 3D**), consistent with the effective suppression of synaptic activity to the recorded neuron. The reduction in local network activity was also reflected in the neuron’s R_m_, which was higher in NBQX, again consistent with decreased synaptic activity (Aw.: 50.6 ± 6.4 MΩ, *n* = 13 neurons, 10 mice vs. +NBQX: 136.2 ± 11.9 MΩ, *n* = 10 neurons, 7 mice, *p* < 0.0001, *unpaired t-test*).

**Figure 3:**
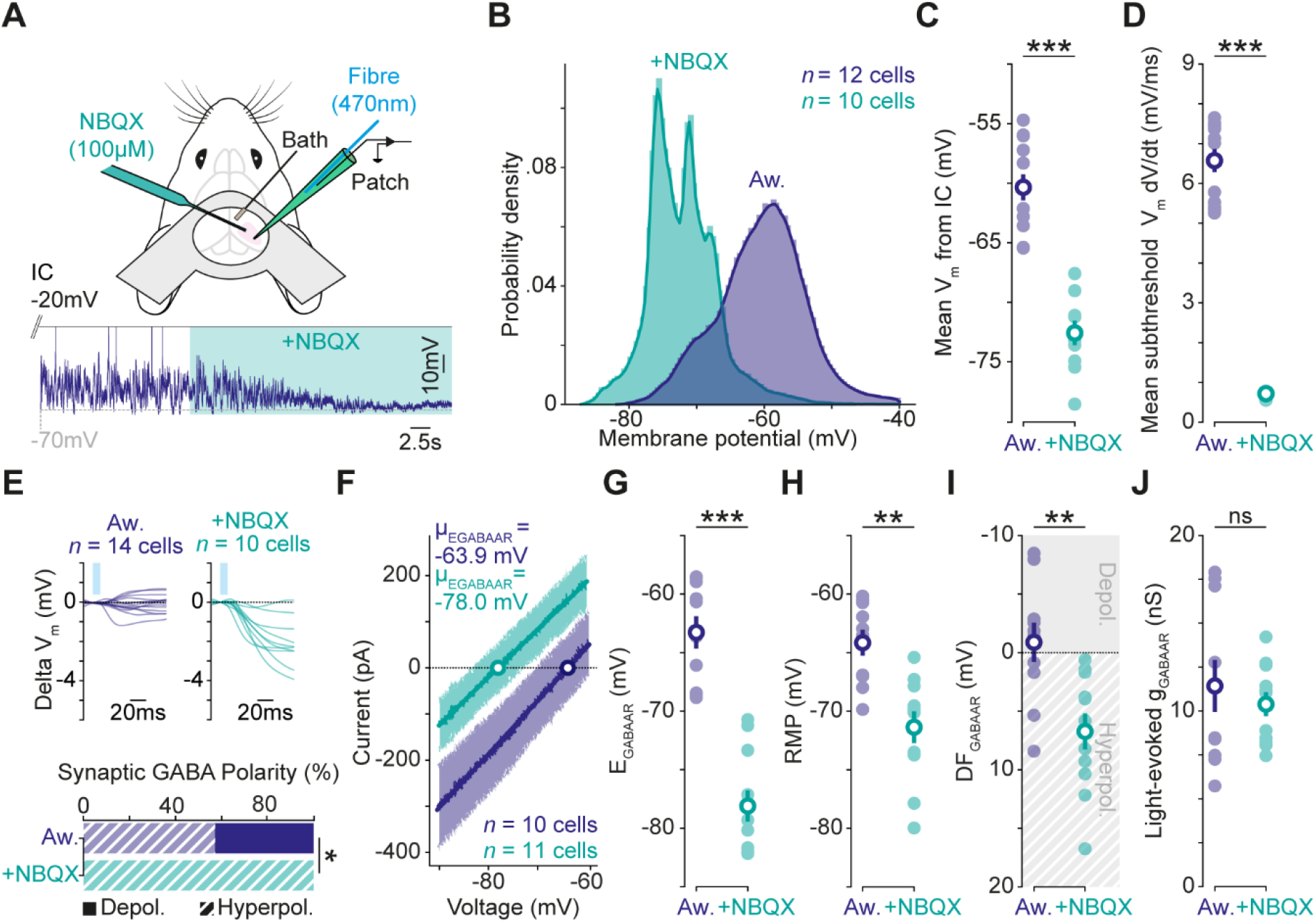
High network activity raises synaptic EGABAAR and promotes shunting fast synaptic inhibition in awake cortex. **(A)** Setup (top) and IC recording (bottom) showing the effects of reducing local network activity by acute, local delivery of NBQX (100μM) in awake cortex. (**B**) Probability density function for V_m_ in control awake cortex and following NBQX (Aw. blue: *n* = 12 neurons, 10 mice vs. +NBQX, turquoise: *n* = 10 neurons, 7 mice). Control data from Figure 2. (**C**) Reducing local network activity caused a hyperpolarizing shift in mean V_m_ (Aw.: -60.3 ± 1.1 mV vs. +NBQX: -72.6 ± 1.0 mV; *p* < 0.0001, *unpaired t-test*). (**D**) Reducing local network activity caused a decrease in the mean change in subthreshold V_m_ (Aw: 6.6 ± 0.3 mV/ms vs. +NBQX: 0.7 ± 0.1 mV/ms; *p* < 0.0001, *unpaired t-test*). (**E**) Averaged light-evoked postsynaptic IC responses (top; Aw., blue: *n* = 14 neurons, 11 mice vs. +NBQX, turquoise: *n* = 10 neurons, 7 mice). Reducing local network activity caused a hyperpolarizing shift in the polarity of light-evoked GABA currents (bottom; Aw.: Depol. 6/14 vs. +NBQX: Depol. 0/10; *p* = 0.03, *Fisher-Exact test*). (**F**) Summary IV plots of light-evoked GABA currents reveal more hyperpolarized synaptic E_GABAAR_ values following local NBQX (Aw.: *n* = 10 neurons, 9 mice vs. +NBQX: *n* = 11 neurons, 7 mice). (**G**) Reducing local network activity led to more hyperpolarized synaptic E_GABAAR_ (Aw: -63.3 ± 1.4 mV vs. +NBQX: -78.1 ± 1.3 mV; *p* < 0.0001, *unpaired t-test*). (**H**) Reducing local network activity caused a hyperpolarizing shift in RMP (Aw.: -64.2 ± 1.1 mV vs. +NBQX: -71.4 ± 1.4 mV; *p* = 0.0006, *unpaired t-test*). (**I**) Reducing local network activity caused a hyperpolarizing shift in DF_GABAAR_ (Aw.: -0.9 ± 1.7 mV vs. +NBQX: 6.7 ± 1.5 mV; *p* = 0.003, *unpaired t-test*). DF_GABAAR_ became different from zero (Aw.: *p* = 0.61, *one-sample t-test*; +NBQX: *p* = 0.001, *one-sample-test*). (**J**) Reducing local network activity did not affect light-evoked synaptic GABA conductances (Aw: 11.4 ± 1.5 nS vs. +NBQX: 10.4 ± 0.7 nS; *p* = 0.52, *unpaired t-test*). ‘ns’, non-significant; **, *p* < 0.01; ***, *p* < 0.001.

To investigate whether the levels of local network activity determine the nature of GABAergic signaling in the awake cortex, the polarity of synaptic GABA_A_R responses was compared across recordings performed with and without NBQX (**Figure 3E**). In the quietened network state, light-evoked responses recorded in current clamp were found to be exclusively hyperpolarizing, which differed to the active awake state (**Figure 3E**). Consistent with this, reducing local network activity caused a hyperpolarizing shift in synaptic E_GABAAR_ (**Figure 3F** and **3G**). When combined with the negative shift in RMP (**Figure 3H**), the net effect of reducing local network activity was to cause a hyperpolarizing shift in DF_GABAAR_, such that the synaptic DF_GABAAR_ was now strongly hyperpolarizing (**Figure 3I**). Importantly, the effects of reducing local network activity with NBQX were not related to effects on the optogenetic paradigm itself, as the amplitude of the light-evoked synaptic GABA_A_R conductances were unaffected by the AMPA receptor antagonist (**Figure 3J**). More generally, this confirmed that the measurements of synaptic E_GABAAR_ were not contaminated by glutamatergic conductances. Also, the differences in synaptic E_GABAAR_ could not be explained by differences in intrinsic membrane properties associated with reducing local network activity (**Figure S3**).

### Shunting inhibition serves to decouple active cortical networks and improve stimulus discrimination

To study the implications of our experimental observations, we constructed simple neural networks that consisted of recurrently connected excitatory pyramidal neurons and inhibitory interneurons (**Figure 4A**). This enabled us to vary synaptic E_GABAAR_ and investigate how an active network responds to different types of input, depending upon whether it is using shunting or hyperpolarizing inhibition. The levels of activity in the networks were maintained by adjusting spike thresholds (**Figure S4**). First, we examined responses to rhythmic input that consisted of sinusoidal waves of a particular frequency (**Figure 4B**). In networks using hyperpolarizing E_GABAAR_, the spiking responses of the excitatory neurons were closely time-locked to the phase of the oscillation and to the other neurons within the network (**Figure 4C**). By contrast, spiking in networks using shunting E_GABAAR_ was more variable with respect to the stimulus phase. To quantify this, we measured the degree of population coupling, which relates the firing of a single neuron to the activity of the surrounding network^27^. This revealed that population coupling was lower in networks that used shunting E_GABAAR_ (**Figure 4D**).

**Figure 4:**
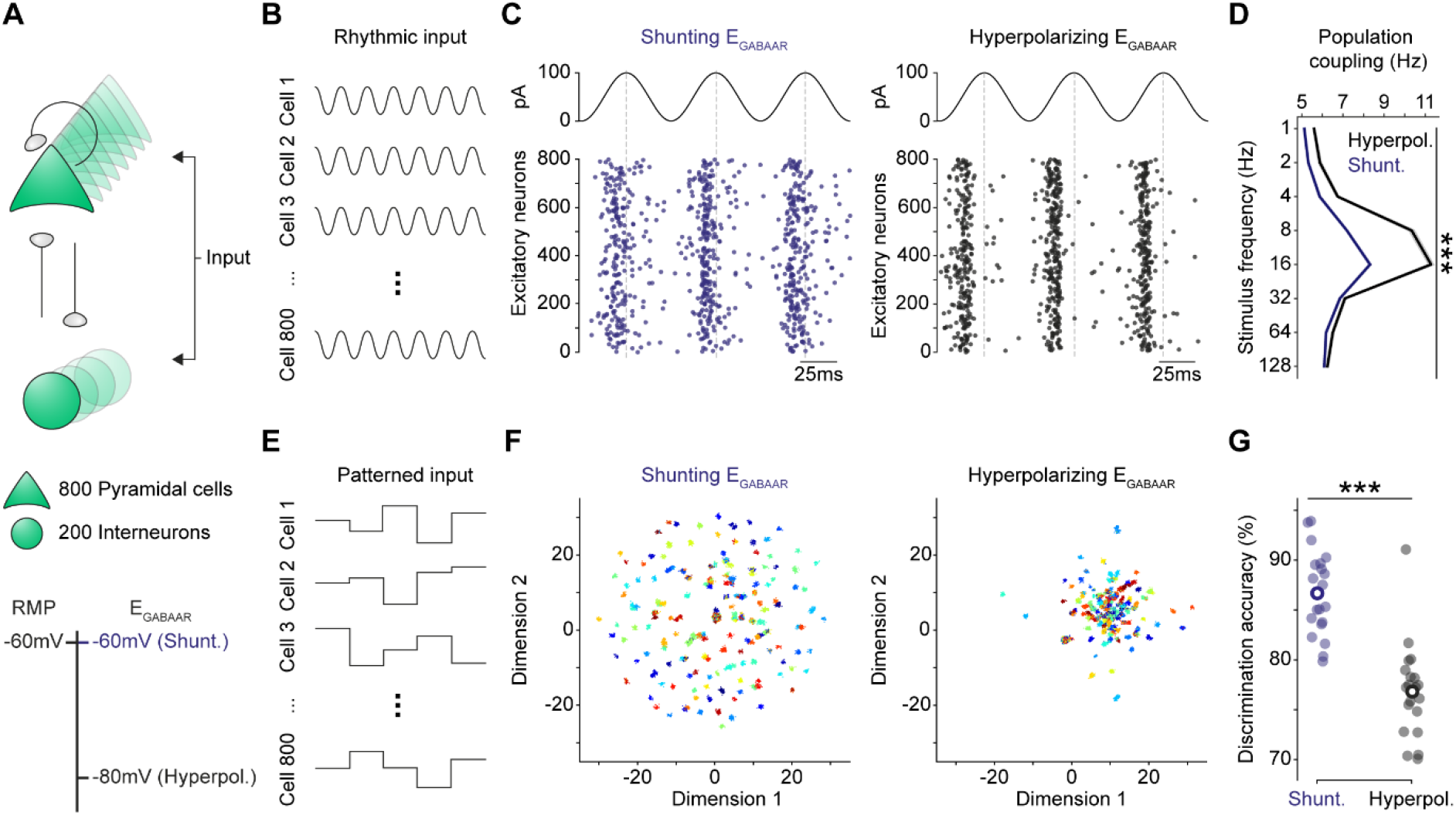
Shunting synaptic EGABAAR serves to decouple active networks and improve stimulus discrimination. **(A)** Schematic (top) illustrating simple network model of recurrently connected glutamatergic and GABAergic neurons. E_GABAAR_ in the glutamatergic neurons (bottom) was set to either equal RMP, consistent with the shunting effects observed active awake cortex (Shunt., blue), or was set to be below the RMP, such that GABA_A_R responses were hyperpolarizing (Hyperpol., black). Activity levels of the network were maintained by adjusting spike threshold in the glutamatergic neurons. (**B**) To investigate population coupling, sine waves of a particular frequency were used to provide the same rhythmic input to all neurons within the network. (**C**) Raster plot showing spiking responses to rhythmic input when the glutamatergic neurons had a shunting E_GABAAR_ (blue, left) or a hyperpolarizing E_GABAAR_ (black, right). (**D**) Population coupling was greater for active networks with hyperpolarizing synaptic E_GABAAR_ (*p* < 0.0001, *two-way ANOVA test with Šídák’s test for multiple comparisons*). (**E**) To explore the effect of synaptic E_GABAAR_ upon stimulus discrimination, patterned input was delivered to the network such that each neuron received a different input value for each stimulus. (**F**) UMAP projections of the neuronal responses to different input patterns. Each dot corresponds to the population vector of spike counts in response to a single input. Each color indicates the different random input patterns. In both conditions, responses to the same input pattern form tight clusters. However, shunting E_GABAAR_ results in improved separability of the responses to different inputs. (**G**) Discrimination accuracy was greater in networks with shunting synaptic E_GABAAR_, than in networks with hyperpolarizing E_GABAAR_ (Shunt.: 86.7 ± 1.1 % vs. Hyperpol.: 76.8 ± 1.1 %, *n* = 20 simulations; *p* < 0.0001, *unpaired t-test*). ***, *p* < 0.001.

Cortical neurons that decouple from the local network tend to respond more independently to incoming stimuli^28,29^. Shunting E_GABAAR_ could, therefore, improve a network’s ability to discriminate between different types of input. To test this idea, we presented patterned inputs and compared the ability of the networks to discriminate between different stimuli (**Figure 4E**). This revealed that networks using shunting E_GABAAR_ were more accurate at distinguishing stimuli (**Figure 4F** and **4G**). Thus, deploying shunting inhibition rather than hyperpolarizing inhibition, can facilitate network decoupling and enhance an active network’s ability to encode stimulus properties.

## Discussion

To determine how fast synaptic inhibition operates *in vivo*, we require measurements that preserve the ionic driving forces acting upon GABA_A_Rs, whilst providing information that can relate a neuron’s synaptic GABA_A_R conductances, synaptic E_GABAAR_, and membrane potential. Whole-cell patch clamp recordings have revealed prominent GABA_A_R synaptic conductances in awake cortex, but compromise the ionic gradients underlying synaptic GABA_A_R transmission^3^. Imaging approaches provide estimates of somatic chloride levels *in vivo*^30,31^, but are limited in their ability to describe synaptic signaling. Also, efforts have been made to infer synaptic GABA_A_R function from whole-cell recordings^21^ or extracellular recordings^32^, although these approaches do not provide direct measurements of E_GABAAR_ or DF_GABAAR_.

By establishing *in vivo* gramicidin perforated recordings and combining these with optogenetic activation of synaptic GABA_A_Rs, we have measured synaptic DF_GABAAR_ in the intact brain. Our recordings demonstrate that the nature of GABA_A_R signaling is linked to the network’s activity. Synaptic GABA_A_Rs in awake cortex exhibit relatively depolarized E_GABAAR_ and low DF_GABAAR_, such that their inhibitory effects are more likely to result from local changes in membrane resistance (i.e. shunting effects). This provides experimental support for theoretical predictions regarding how E_GABAAR_ reflects a dynamic equilibrium between Cl^-^ extrusion and intrusion processes, and how Cl^-^ influxes associated with high GABA_A_R activity can increase the Cl^-^ load experienced by a neuron^8^. Our observations have implications for how fast synaptic inhibition might vary under other *in vivo* network conditions that are associated with changes in ongoing synaptic activity^33,23^, and how such short-term, activity-dependent changes may interact with longer-term fluctuations in Cl^-^ homeostasis^34^. In addition, the space-clamp limitations associated with *in vivo* patch-clamp recordings^35,36^ mean that our measurements are most relevant for somatic GABA_A_R synapses and future work could explore effects upon GABA_A_R synapses in different neuronal compartments.

A synaptic E_GABAAR_ that favors shunting has important implications for neural computation. This includes effects upon synaptic integration^12^, gain modulation^13^ and network synchronization^37^. In the case of synaptic inhibition that is purely shunting, the duration of inhibition is restricted to the time course of the GABAAR conductance. By contrast, the effects of hyperpolarizing GABAAR-mediated inhibition are more long-lasting, such that they can synchronize the recovery of neurons within the network^38^. Our modelling experiments indicate that the shunting synaptic E_GABAAR_ observed in active awake cortical networks favors a functional decoupling of neurons, which can enhance stimulus discrimination, reminiscent of ‘soloist’ neurons that show weak coupling to the network^28,29^. Our findings therefore suggest that GABA_A_R signaling adapts to optimize cortical function, and that the cortex can use dynamic changes in GABA_A_R-mediated synaptic inhibition to regulate the functional coupling of the network.

## Methods

### Experimental model

All mice were bred, housed and used in accordance with the United Kingdom Animals (Scientific Procedures) Act (1986). Homozygous Gad2^tm2(Cre)Zjh/J^ mice (Gad2-IRES-Cre) were crossed with homozygous B6;129S-Gt(ROSA)^26Sortm32(CAG-COP4*H134R/EYFP)Hze/J^ mice (Ai32). This produced a heterozygous colony expressing channelrhodopsin-2 (ChR2(H134R)-YFP) in Gad2-positive neurons, which includes the main subclasses of GABAergic interneurons^20^. Mice were purchased from Jackson Laboratory (Maine, USA). Both male and female mice were used in the experiments. Mice were maintained under a 12-hour:12-hour light-dark cycle and fed *ad libitum*. For all experiments both males and females were used and mice were between six to eight weeks postnatal age at the time of recording.

### Surgical procedures

The preparation for anaesthetized recordings was adapted from previously published protocols^39–41^. Mice were anaesthetized with an intraperitoneal (IP) injection of 25 % urethane (1 g/kg, diluted in sterile PBS). To counteract adverse events caused by urethane, a bolus of the anticholinergic agent, Glycopyrronium Bromide (0.01 mg/kg) was administered subcutaneously (SC). Local anesthetic (Marcain 2 mg/kg) was applied intradermally to the scalp and topically in the ears prior to mounting the mouse into the head holding apparatus (Narishige) under a surgical stereoscope (Olympus). The mouse’s body temperature was maintained at 37 degrees Celsius using a heating mat and rectal probe. The animal’s head was shaved and eye-protecting ointment (Viscotears) was applied to both eyes. An incision in the scalp was made using surgical scissors and the area expanded with blunt dissection to expose the skull. The site of the craniotomy was marked over the primary visual cortex (V1). Tissue adhesive (Vetbond) was applied to fix the surrounding scalp to the skull and to secure cranial sutures. Multiple layers of dental cement (Simplex Rapid) were applied to create a recording chamber on top of the skull. A 0.5 mm craniotomy was drilled over the marked region using a dental drill (Foredom). The craniotomy was submerged in cortex buffer (containing, in mM: 125 NaCl, 5 KCl, 10 HEPES, 2 MgSO_4_·7H_2_O, 2 CaCl_2_·2H_2_O, 10 Glucose). The bone flap and dura were removed. The animal was then transferred to the *in vivo* patch setup, and the recording session typically lasted 3 hours between zeitgeber time 3 (ZT3) and ZT6, at which point the animal was culled.

The preparation for awake recordings consisted of three phases based on published protocols^39,42^. The first phase involved the fixation of the head plate. Mice were anaesthetized with isoflurane (Zoetis) and mounted into a stereotaxic frame (Kopf). Subcutaneous analgesia (meloxicam 5mg/kg and buprenorphine 0.1mg/kg) was administered along with intradermal local analgesic (marcain 2 mg/kg) into the scalp. The scalp was shaved (Wahl) and cleaned (Hibiscrub). Eye-protecting ointment (Viscotears) was applied. The scalp was then removed and the site was washed with sterile cortex buffer (containing, in mM: 125 NaCl, 5 KCl, 10 HEPES, 2 MgSO_4_·7H_2_O, 2 CaCl_2_·2H_2_O, 10 Glucose). After drying the skull with adsorbent swabs (Haag-Streit), the periosteum was removed using a micro curette (Fine Science Tools). Tissue adhesive (Vetbond) was applied to secure cranial sutures and to fix the surrounding scalp to the underlying bone. A custom-designed aluminum headplate with a 7 mm well was bonded to the skull, first with adhesive glue (Loctite), and then followed by serial layers of dental cement (Super-Bond). The well was then covered with silicone sealant (Kwik-Cast). The animal was singly housed and allowed to recover. From day three following head plate fixation, the animal was habituated to head-fixation for increasing time intervals up to 60 minutes. On the day of the recording, the animal was briefly anaesthetized with isoflurane and mounted onto a stereotaxic frame. The silicone sealant was removed and the area washed with sterile cortex buffer. A 0.5 mm craniotomy was created using a dental drill (Foredom) and the bone flap removed. A durectomy was performed and the site was covered with a soft dressing soaked in cortex buffer. This step in the procedure was limited to 20 minutes. The animal was then remounted onto the head-fixation setup and transferred to the *in vivo* patch setup. The mouse was allowed to fully recover for at least 30 minutes before recording was commenced. Recording sessions typically lasted 2-3 hours between ZT3 and ZT6, at which point the animal was culled.

### Electrophysiological recordings

All electrophysiological recordings were performed using an Axopatch 700A amplifier with data acquired at 20 kHz using a Digitimer 1550 system controlled by Clampex software (Molecular Devices). A HumBug noise eliminator (Digitimer) was used to remove 50 Hz noise. To perform perforated patch-clamp recordings, the internal pipette solution was prepared immediately prior to recording by combining a high chloride (150 mM) solution (in mM: 141 KCl, 9 NaCl, 10 HEPES) heated to 37 degrees Celsius, with a stock solution of gramicidin A (4 mg/ml - dissolved in dimethyl sulfoxide, DMSO, Merck) to achieve a final concentration of 80 μg/mL gramicidin^19^. The solution was then vortexed (40 seconds) and sonicated (20 seconds). The patch pipette was back-filled with the gramicidin solution and mounted on a Optopatcher pipette holder (A-M Systems) which contained a 50 μm fiber (Thorlabs) connected to a 473nm laser (MBL-FN-473-150mW, CNI Laser). Pipettes were lowered onto the brain surface and blind patching commenced. Once the gigaseal had formed, perforation was then monitored by observing changes in series resistance. Recording protocols were started once the series resistance had stabilized at <100 MΩ. Rupture or breakthrough of the perforation in to whole-cell configuration was detected by a sudden and persistent depolarization of the equilibrium potential of the GABA_A_R (E_GABAAR_), consistent with dialysis of the neuron with the high chloride pipette solution. In a subset of experiments, KCC2 was blocked by injecting the selective antagonist, VU0463271^43^, directly into the cortex. The injection pipette contained 100 μM VU0463271 (Tocris) in ACSF, which was delivered at a rate of 33 nL/min to a total volume of 200 nL. In a subset of experiments where local network activity was reduced, the AMPA receptor blocker 2,3-dihydroxy-6-nitro-7-sulfamoylbenzo (F) quinoxaline (NBQX) was injected directly into the cortex^44^. The injection pipette contained 100 μM NBQX (Tocris) in ACSF, which was delivered at a rate of 33 nL/min to a total volume of 250 nL.

### Recording protocols and data analysis

Data was acquired using recording protocols configured in Clampex (Molecular Devices) and analyzed using custom code written in Python. Online series resistance compensation was not used, as the high amounts of activity in the *in vivo* brain would cause large fluctuations in input current, which increases the rate of perforation rupture^45,46^. Therefore, to correct for series resistance effects, offline correction was performed^47^. The voltage drop caused by the series resistance (R_s_) was calculated by multiplying the measured current response with 90 % of the R_s_. The voltage drop was then subtracted from the command voltage to estimate the neuron’s membrane potential. Membrane and recording properties were calculated by measuring the change in current in response to a -10 mV step during voltage-clamp recordings. R_s_ was calculated from both the peak current elicited by the -10 mV voltage step, and by estimating the peak after fitting an exponential to the decay of the current transient response to the -10 mV step. These methods gave similar values and so the numerical average was used as a final estimate of R_s_. To calculate the membrane resistance (R_m_), R_s_ was subtracted from the measured input resistance.

To determine the resting membrane potential (RMP) during current-clamp recordings, spontaneous activity was recorded for a continuous period of five minutes. During analysis, membrane potential values greater than -40 mV were removed to avoid distortions caused by action potentials. Average distribution plots were created by concatenating all membrane potential values across all neurons in each group, to which a Gaussian kernel-density estimate was then fitted. The level of subthreshold synaptic activity was determined by measuring the rate of change in the membrane potential over time, after excluding action potentials. The rate of change in membrane potential was calculated by taking the first derivative (V_m_ dV/dt in mV/ms). The first derivative was winsorized and the mean of the modulus of the V_m_ dV/dt calculated for each neuron. Synaptic GABA_A_R-mediated responses in current clamp were evoked with a 10 ms light pulse, presented during a one second sweep. The average of 15 sweeps was then calculated and normalized to the mean membrane potential during the 100 ms preceding the light pulse. Current-clamp recordings from a subset of the neurons in the NBQX condition also contributed to another study^34^. The polarity of the synaptic GABA_A_R response was defined as the mean normalized membrane potential during the 100 ms following the light pulse. If the mean value was greater than zero, the response was classified as depolarizing, whereas a mean value below zero was classified as hyperpolarizing.

Two types of voltage-clamp protocol were used to measure synaptic E_GABAAR_: a ramp protocol and a step protocol. The ramp protocol involved clamping the neuron at -70 mV, and then imposing two consecutive voltage ramps, each lasting 150 ms, which extended from 60 mV below the holding voltage to 40 mV above the holding voltage (i.e. from -130 mV to -30 mV, at a rate of 0.7 mV/ms). The first ramp (i.e. the ‘baseline’ ramp) sampled the neuron’s intrinsic membrane currents and the second ramp (i.e. the ‘light’ ramp) included a light-evoked synaptic GABAAR conductance. The light-evoked synaptic GABAAR conductance was elicited with a 10 ms light pulse that was initiated 30 ms prior to the start of the ramp, to ensure the evoked GABAAR current was at its peak during the ramp^22^. During analysis, the first and last 15 ms of each ramp were excluded to avoid transient currents caused by the discharge of the pipette capacitance. The currents from both ramps were then superimposed on a current-voltage (IV) plot using the series corrected membrane potential and, to avoid action potentials and capacitance transients, the current responses were cropped to only include regions that were clear of these sources of contamination. A straight line was fitted to both currents. The current from the baseline ramp was used to infer the equilibrium potential of the holding current (from which the resting membrane potential, RMP, could be inferred), defined as the voltage at which the fitted line was equal to zero. The point at which the fitted lines for the two ramps intersected is E_GABAAR_. This can also be calculated by subtracting the current response during the baseline ramp, from the current response during the light ramp. The voltage at which a fitted line to this subtracted current is equal to zero, is equivalent to E_GABAAR_. The driving force can then be calculated by subtracting the measured E_GABAAR_ from the RMP. The slope of the subtracted current is equal to the GABA_A_R conductance. The step protocol meanwhile, involved estimating synaptic E_GABAAR_ from a voltage-clamp protocol in which the neuron was exposed to a series of voltage steps, whilst eliciting a light-evoked GABA response during each step. The voltage was stepped in 10 mV increments from -130 mV to -30 mV. Each step lasted 500 ms, with a 10 s interval between steps. The light-evoked synaptic GABA conductance was evoked 100 ms after the start of each voltage step, by delivering a 100 ms light pulse. For analysis purposes, the membrane current was measured immediately before the light pulse (i.e. ‘baseline’ current) and then 30 ms after the onset of the light pulse, which corresponds to the peak of the GABA_A_R conductance (i.e. ‘light’ current). The two currents were then analyzed in a similar way to that described for the ramp protocol.

### Neuronal network simulations

Network models were used to explore the effect of different synaptic E_GABAAR_ upon population coupling and stimulus discrimination by an active cortical network. The network was constructed using the neuron simulator Brian 2^48^ and comprised 800 glutamatergic neurons and 200 GABAergic neurons. Each neuron was modelled as a single compartment, current-based leaky integrate-and-fire neuron. Free parameters were set as shown in **Table 1**. The glutamatergic neurons received synaptic connections from both the glutamatergic and GABAergic neurons; the GABAergic neurons only received connections from the glutamatergic neurons. Within these constraints, the connection probability was set uniformly to 10 %. Glutamatergic synaptic weights were set to 0.3-1 nS and GABAergic synaptic weights were initialized at 1-5 nS. To explore the effect of synaptic E_GABAAR_ upon population coupling we provided a sinusoidal input current to all neurons oscillating between 0 and 105 pA. To prevent full synchronization of all neurons, each neuron also received a random, independent input current with values drawn from a normal distribution with mean and standard deviation of 52.5 pA, varied every 10 ms. In other words, 50% of the total external input to each neuron was synchronous and 50 % was asynchronous. To explore the effect of synaptic EGABAAR upon stimulus discrimination, we provided patterned input to the network. Each pattern consisted of 1000 random input currents chosen uniformly between 0 and 105 pA, one for each neuron. Each pattern was presented to the network 100 times, in random order, and each presentation had a duration of 100 ms. To determine the separability of the neuronal responses, we computed the spike count for each presentation, then trained and tested a k-nearest neighbor classifier using 5-fold stratified cross-validation. To investigate how shunting inhibition and hyperpolarizing inhibition affect an active network’s responses to different types of input, synaptic E_GABAAR_ in all glutamatergic neurons within a network was set to either -60 mV (‘shunting’) or -80 mV (‘hyperpolarizing’). E_GABAAR_ was set to -60 mV in the GABAergic neurons. The overall activity levels were matched across the different networks by adjusting the spike threshold in the glutamatergic neurons, so as to produce an equivalent number of spikes in simulations with the same stimulus (**Figure S4**). Each simulation was repeated 20 times.

**Table 1:** Free parameters used for simple neuronal network modelling.

|  |  |
| --- | --- |
| Equilibrium potential of the leak current ($E_{\text{Leak}}$, equivalent to RMP) | -60 mV |
| Equilibrium potential of the glutamatergic current | 0 mV |
| Excitatory postsynaptic potential decay time constant | 5 ms |
| Inhibitory postsynaptic potential time constant | 10 ms |
| Membrane capacitance | 200 pF |
| Membrane time constant | 5 ms |
| Refractory period | 5 ms |

### Simulating measurements of synaptic E_GABAAR_

To investigate the effect of intrinsic membrane properties on our estimates of synaptic E_GABAAR_ under different network conditions, a multi-compartment model of an adult pyramidal neuron from L2/3 of mouse primary visual cortex was constructed using the NEURON simulation environment^49^. The neuron’s morphology was sourced from NeuroMorpho^50^ and based on a reconstruction (NMO_62358) that was shared by Madisen and colleagues^51^. Model parameters are shown in **Table 2**.

**Table 2:** Parameters used for modelling effects of membrane resistance.

|  |  |
| --- | --- |
| Membrane capacitance | $C_m = 2.515 \mu\text{F}/\text{cm}^2$ |
| Axial resistance ($R_{\text{axial}}$) | 150 $\Omega\text{cm}$ |
| Passive membrane reversal (equivalent to RMP) | An. $e_{\text{pas}} = -70.7 \text{ mV}$<br>Aw. $e_{\text{pas}} = -64.2 \text{ mV}$<br>+NBQX $e_{\text{pas}} = -71.4 \text{ mV}$ |
| Voltage clamp electrode series resistance ($R_s$) | 47.5 MΩ |
| Passive membrane resistance | An. $R_m = 10^{3.784} \Omega\text{cm}^2$<br>Aw. $R_m = 10^{3.540} \Omega\text{cm}^2$<br>+NBQX $R_m = 10^{3.988} \Omega\text{cm}^2$ |

With these parameters, the membrane resistance (R_m_) measured by simulated voltage clamp at the soma was 87.2 MΩ in the anaesthetized condition, 50.6 MΩ in the awake condition and 136.2 MΩ in the Awake + NBQX condition. These values matched the average experimentally measured membrane resistance from data (**Figure S3**). Activation of GABA_A_Rs was simulated by placing twenty GABA_A_R-containing synapses randomly within a 75 μm radius of the center of the soma. E_GABAAR_ was set to between -85 mV and -35 mV (iterated by 5 mV for each simulation). Activation of GABA_A_R synapses was simulated by using an alpha function with a tau of 150 ms. The peak local conductance of each GABA_A_R synapse was set to 2 nS. To simulate the experimental estimation of synaptic E_GABAAR_, a simulated voltage clamp was placed at the soma. Two consecutive voltage ramps were then applied, one before and one during simulated activation of GABA_A_Rs (to reproduce the experimental protocol). Synaptic E_GABAAR_ was estimated using IV plots, either with 0% R_s_ correction, or 90 % R_s_ correction, to replicate the experimental data acquisition and analysis process.

## Acknowledgements

We would like to thank members of the Akerman lab for advice and comments. We thank Mateo Velez-Fort; Christopher Pugh and Troy Margrie (Sainsbury Wellcome Centre) for their help in setting up the *in vivo* patch-clamp recordings, and Ed Mann for his comments on the manuscript. We thank Lex Kravitz, Luigi Petrucco and Ethan Tyler for sharing their mouse illustrations on the Sci-Draw open-source platform. The research leading to these results has received funding from the European Research Council under grant agreement 617670; plus BBSRC project BB/S007938/1. In addition, this work was supported by a Shaun Johnson Memorial Scholarship sponsored by the Leverhulme Trust and Mandela Rhodes Foundation (R.J.B.) and a Wellcome Trust International Intermediate Fellowship 222968/Z/21/Z to J.V.R.

## Author contributions

Conceptualization: R.J.B.; C.J.A.

Investigation: R.J.B.; P.J.N.B.; J.V.R.

Writing: R.J.B.; C.J.A.

Supervision: C.J.A.; A.S.

Funding acquisition: R.J.B.; C.J.A.; A.S.

## Supplemental information

**Supplementary Figure 1:**
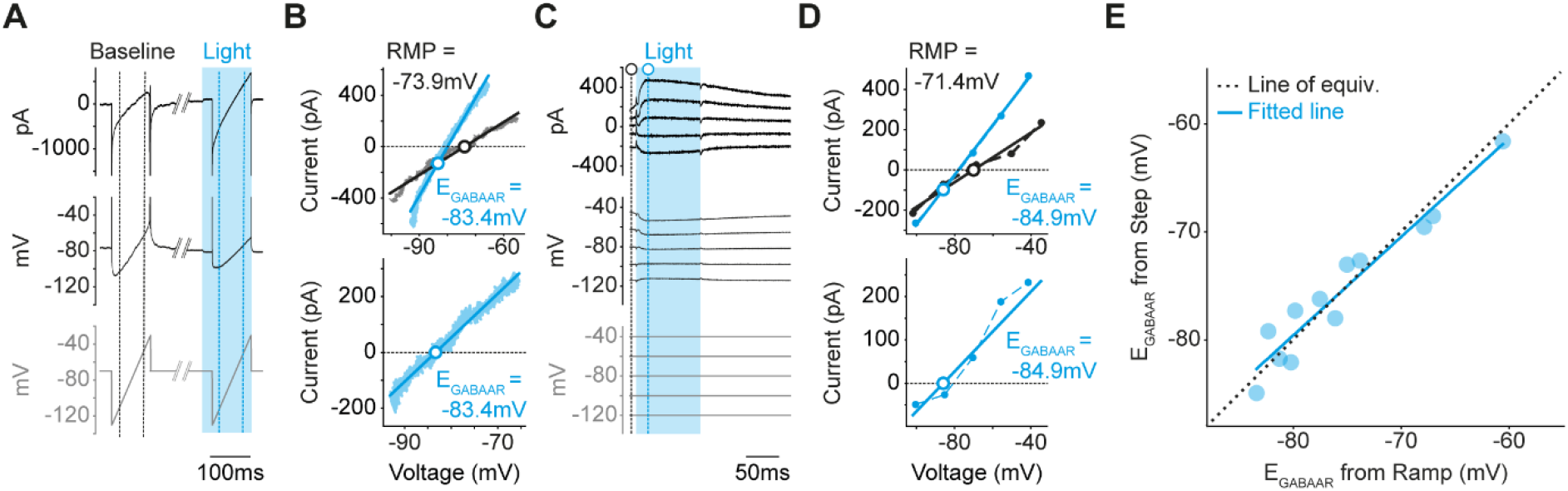
*In vivo* voltage-ramp protocols provide accurate estimates of synaptic E_GABAAR_. **(A)** Raw trace of a ramp protocol showing the command holding voltage (bottom, grey), the voltage after applying series resistance correction (R_s_ = 52.1 MΩ, middle, black), and the current response (black, top). The protocol consisted of a control ramp (‘baseline’) and a second ramp during which the light-evoked postsynaptic GABA_A_R-mediated synaptic current was evoked with a blue light pulse (‘light’; cyan shaded area). The pairs of vertical dashed lines indicate the regions of the ramps that were analyzed. (**B**) IV plots from a ramp protocol (top) in which the baseline (black) and light (cyan) current responses are shown. The intersection of the two fitted lines represents synaptic E_GABAAR_, while the point at which the baseline current crosses zero is equivalent to the neuron’s RMP. IV plot of the subtracted current (bottom), which corresponds to the synaptic GABA_A_R current and has a value of zero at E_GABAAR_. (**C**) A voltage step protocol was used to estimate synaptic E_GABAAR_ in the same neuron as in ‘A’. The voltage step protocol (bottom, grey), the voltage after applying series resistance correction (R_s_ = 50.8 MΩ, middle, black) and current response (top, black) are shown. GABA_A_R-mediated currents were evoked using light pulses (470nm, 100ms, cyan shading). Black vertical dashed line indicates where baseline current was calculated. Cyan vertical dashed line indicates where measurements were made of the GABAAR-mediated synaptic current, 20-30 ms from light onset. Previous work has shown that this isolates the GABAAR response ^22^. (**D**) IV plots (top) of the baseline current (black) and the current during the light-evoked GABA_A_R response (cyan). IV plot of the subtracted current (bottom), which corresponds to the synaptic GABA_A_R current and has a value of zero at E_GABAAR_. (**E**) Scatter plot showing the correlation between E_GABAAR_ calculated from the ramp and voltage step protocols for each neuron (*n* = 12 neurons, 6 mice). A strong correlation was observed between synaptic E_GABAAR_ measurements from the ramp protocol and the GABA_A_R-mediated synaptic current in the step protocol (cyan, *R squared* = 0.92, *p* < 0.0001, *n* = 12 neurons from 6 mice, *Wald test*).

**Supplementary Figure 2:**
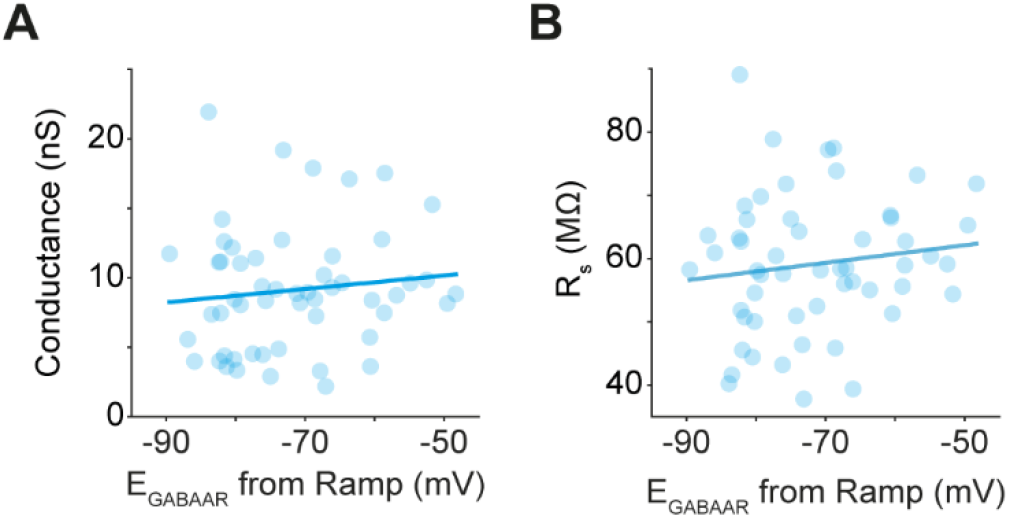
Estimates of synaptic E_GABAAR_ are not related to the amplitude of the light-evoked postsynaptic GABA conductance or the series resistance. **(A)** No correlation was observed between synaptic E_GABAAR_ and the conductance of the light-evoked GABA_A_R response (*pooled mean* 10.62 ± 0.6 nS, *R*^*2*^ = 0.01, *p* = 0.41, *n* = 54 neurons from 39 mice). Conductance was calculated from the slope of the GABA_A_R current recorded during the voltage ramp. (**B**) No correlation was observed between synaptic E_GABAAR_ and the neuron’s series resistance (R_s_, *pooled mean* 59.13 ± 1.5 MΩ, *R*^*2*^ = 0.02, *p* = 0.33).

**Supplementary Figure 3:**
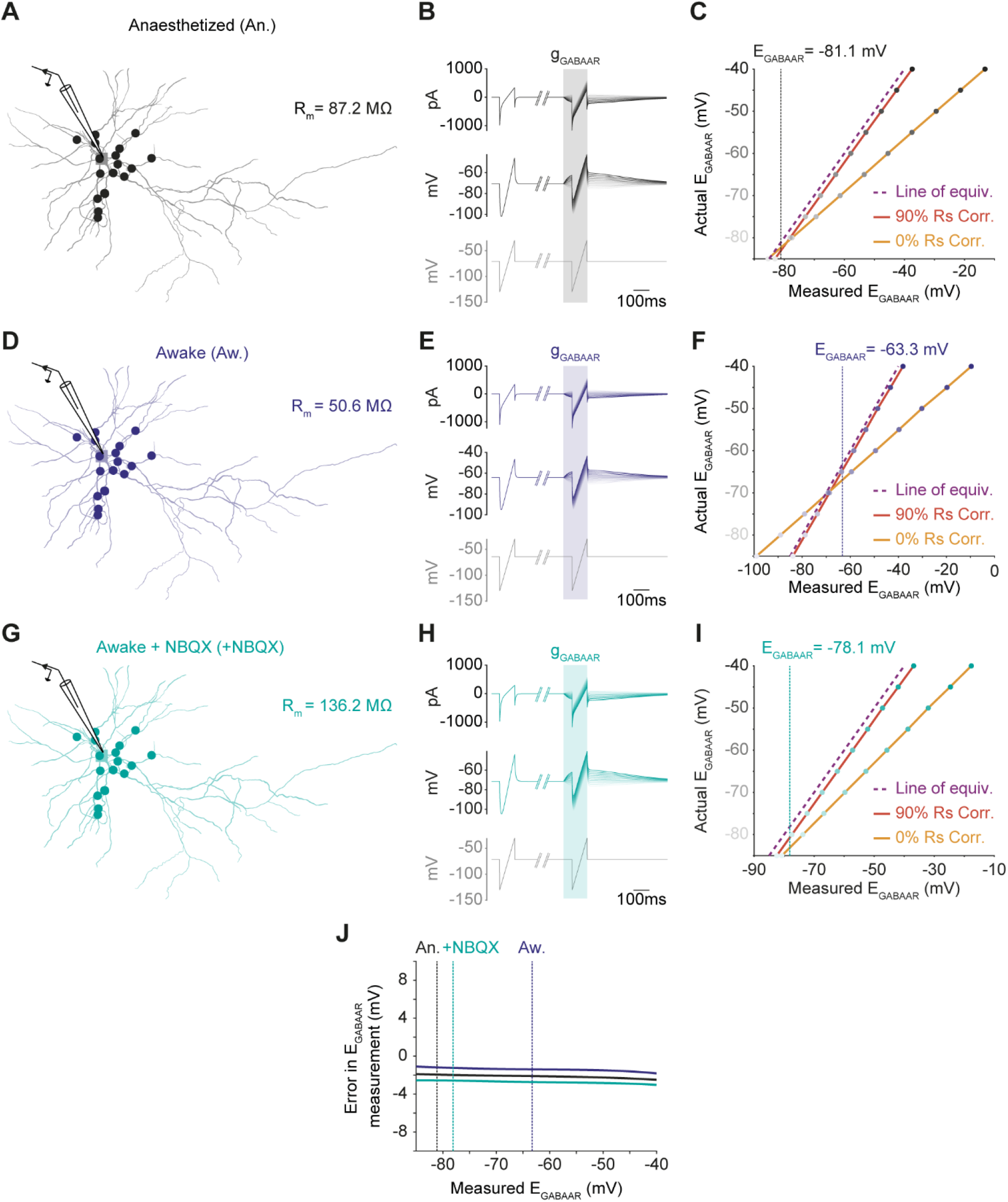
Differences in intrinsic membrane properties have minimal effects upon estimates of synaptic E_GABAAR_. To simulate the effect of intrinsic membrane properties on the estimation of synaptic E_GABAAR_ under different experimental conditions, a multi-compartment model of a L2/3 pyramidal neuron from mouse V1 was constructed using NEURON. (**A**) Morphological reconstruction of the L2/3 pyramidal neuron, with the location of GABA_A_R-mediated synaptic inputs indicated (filled circles). To recapitulate the anaesthetized recording conditions (An., black), the membrane resistance (R_m_) in the model neuron was set to the mean experimentally observed value in our recordings from anaesthetized cortex. (**B**) Voltage-ramp protocols that matched the experimental protocols were applied to the model neuron and used to generate measurements of synaptic E_GABAAR_, which could be compared to the actual E_GABAAR_ that had been preconfigured in the simulated neuron. The different line intensities indicate simulations using different preconfigured E_GABAAR_ values, ranging from -85 mV (lightest line) to -40 mV (darkest line) in 5 mV increments. (**C**) Plot showing the relationship between the actual E_GABAAR_ and measured E_GABAAR_ in the anaesthetized model neuron. The line of equivalence (purple) reflects where the actual and measured E_GABAAR_ values are equal. The relationship between the actual and measured E_GABAAR_ values was determined with zero series resistance correction (‘0% R_s_ corr’, orange) and with 90% series resistance correction, which corresponds to experimental conditions (‘90% R_s_ Corr.’, red). The vertical dashed line indicates the experimentally observed E_GABAAR_ in the anaesthetized state. (**D**) As in ‘A’, except that to recapitulate the awake state (Aw., blue) the membrane resistance and RMP in the model neuron was set to the mean experimentally observed values in our recordings from awake cortex. (**E**) As in ‘B’, showing simulations of the awake state. (**F**) As in ‘C’, with the vertical dashed line indicating the experimentally observed E_GABAAR_ in the awake state. (**G**) As in ‘A’, except that to recapitulate the awake state under conditions of reduced local network activity (+NBQX, turquoise) the membrane resistance and RMP in the model neuron was set to the mean experimentally observed values in our recordings from awake cortex plus NBQX. (**H**) As in ‘B’, showing simulations of the awake cortex plus NBQX state. (**I**) As in ‘C’, with the vertical dashed line indicating the experimentally observed E_GABAAR_ in the awake cortex plus NBQX state. (**J**) The estimated error in E_GABAAR_ measurements (Actual E_GABAAR_ – Measured E_GABAAR_) plotted for each of the three cortical states (using the 90% R_s_ corrected data). The experimentally measured E_GABAAR_ for each state is indicated with the vertical dashed lines. Across the three states, the estimated error for measuring synaptic E_GABAAR_ was 1-3 mV.

**Supplementary Figure 4:**
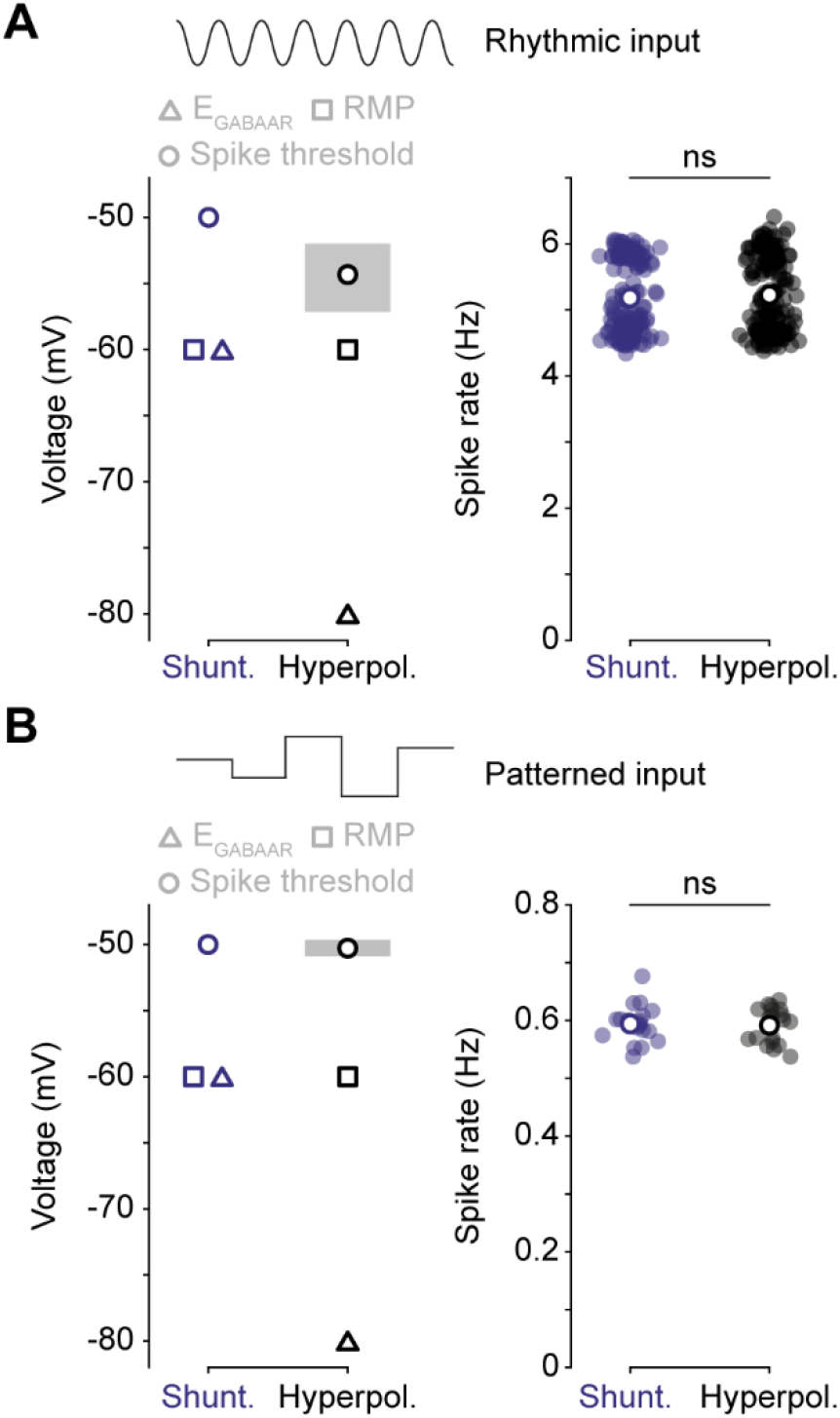
Comparing active networks that utilize different modes of synaptic inhibition. Neural networks were configured to compare the effects of shunting inhibition (E_GABAAR_ = 60mV) and hyperpolarizing inhibition (E_GABAAR_ = -80 mV) on processing different types of input stimuli. The resting membrane potential (RMP) was set to -60 mV in all networks. The spike threshold was set to -50 mV in the shunting network and was allowed to vary in the hyperpolarizing network, so as to achieve comparable levels of overall spiking activity. (**A**) Schematic (left) showing the parameters used in networks that received rhythmic input stimuli. The E_GABAAR_ (triangle), RMP (square) and spike threshold (circle) are indicated. The shaded region illustrates the spike threshold values in hyperpolarizing networks (mean: -54.23 mV). The spike rate (right) was confirmed as equivalent between the shunting and hyperpolarizing networks during presentation of the rhythmic input frequencies (Shunt.: 5.18 ± 0.04 mV vs. Hyperpol.: 5.22 ± 0.05 mV, *n* = 160, *p* = 0.49, *unpaired t-test*). (**B**) Schematic (left) showing the parameters used in networks that received patterned input stimuli. Shaded region illustrates the spike threshold values in hyperpolarizing networks (mean: -50.25 mV). The spike rate (right) was confirmed as equivalent between the shunting and hyperpolarizing networks during presentation of the patterned input (Shunt.: 0.594 ± 0.007 mV vs. Hyperpol.: 0.591 ± 0.007 mV, *n* = 20, *p* = 0.81, *unpaired t-test*). ‘ns’, non-significant.

